# Arabidopsis BAG proteins regulate cellulose synthase stability

**DOI:** 10.64898/2026.09.12.751142

**Authors:** Peng Wang, Isabelle Vanhoutte, Lise Charlotte Marie Noack, Evelien Mylle, Neeltje Schilling, Wei Siao, Xiaohong Zhuang, Yasin Dagdas, Thomas B. Jacobs, Staffan Persson, Daniel Van Damme, Eugenia Russinova

## Abstract

Cellulose synthase complexes (CSCs) synthesize cellulose at the plasma membrane, and their activity and trafficking are critical for maintaining cell wall integrity during plant growth. Clathrin-mediated endocytosis (CME) regulates CSC internalization and has been implicated in their rapid stress-induced removal from the plasma membrane. Stress adaptation, instead, requires the maintenance of a subset of CSCs at the plasma membrane, yet the mechanisms underlying this homeostasis remain poorly understood. The Arabidopsis Bcl-2-associated athanogene4 (BAG4) was identified as an interactor of the adaptor protein 2 complex (AP-2) and the TPLATE complex (TPC), two key components of plant CME. Here, we show that AP-2 and the TPC associated with four closely related BAG proteins, BAG1-BAG4. A quadruple mutant exhibited abnormal growth, increased sensitivity to salt stress, and reduced endocytic flux. However, the abundance, localization and dynamics of CME machinery was largely unaffected, suggesting that BAG proteins are not core regulators of CME. Instead, BAG1-BAG4 deficiency caused hypersensitivity to cellulose biosynthesis inhibitors and impaired hypocotyl elongation in darkness, consistent with defective cellulose-dependent growth. BAG1-BAG4 also interacted with CESA6, and salt-induced CESA6 degradation and ubiquitination was enhanced in the quadruple mutant. Together, these findings identify BAG1-BAG4 as redundant proteostasis factors that safeguard CESA6 stability during salt stress, thereby maintaining cellulose synthesis, cell wall integrity, and plant stress tolerance.

**Significance Statement:** Cellulose synthase complexes produce cellulose at the plasma membrane, but how plants maintain these complexes under environmental stress remains unclear. Here, we identify the Arabidopsis BAG family proteins BAG1-BAG4 as redundant proteostasis regulators that safeguard the primary cellulose biosynthesis machinery during salt stress. Although BAG1-BAG4 associate with key components of the endocytic machinery, they do not appear to function as core regulators of clathrin-mediated endocytosis. Instead, they limit stress-induced ubiquitination and internalization of cellulose synthases, thereby maintaining their abundance at the plasma membrane.

## Introduction

The plant cell wall, consisting primarily of cellulose, pectins, hemicelluloses, lignin, and structural proteins, is crucial for growth and environmental adaptation (1–3). Hemicelluloses, pectins, and glycoproteins are synthesized in the Golgi apparatus and transported to the apoplast via exocytosis (4). In contrast, cellulose is synthesized directly at the plasma membrane (PM) by CELLULOSE SYNTHASE (CESA) complexes (CSCs) (5). Like other PM-localized proteins, CSCs are synthesized in the endoplasmic reticulum and trafficked through the Golgi and *trans*-Golgi network (TGN) to the PM. Small CESA compartments (SmaCCs), also known as microtubule-associated CESA compartments (MASCs), act as transient trafficking intermediates that facilitate CSC delivery to and recycling from the PM (6).

Previous studies have demonstrated that CSCs undergo clathrin-mediated endocytosis (CME) (7–9). CME is an evolutionarily conserved internalization pathway that depends on a network of proteins, including the cage-forming protein clathrin, adaptor proteins, and accessory proteins, which work together to select and enclose cargo within the assembled clathrin-coated vesicles (CCVs) (10–16). CME regulates the PM abundance of diverse proteins, including receptor kinases and transporters (17–21). In plants, this process is mediated by AP-2 and the TPLATE complex (TPC), two core adaptor protein complexes (22). AP-2 and TPC are also required for the internalization of CSCs (7–9). Salt and osmotic stresses induce a rapid depletion of CSCs from the PM and their accumulation in SmaCCs/MASCs, while CME has been implicated in CSC internalization and stress-induced SmaCC/MASC formation (23, 24).

Bcl-2-associated athanogene4 (BAG4) was identified as an accessory protein for AP-2 and TPC, as it interacts with multiple subunits of both adaptor complexes (15). This finding suggests a possible role for BAG4 in CME. BAG4 belongs to the evolutionarily conserved BAG protein family, which is characterized by the presence of a C-terminal BAG domain (25–27). In mammals, this domain mediates interactions with the heat shock protein70/ heat shock cognate protein70 (HSP70/HSC70) chaperones, enabling BAG proteins to participate in various signaling pathways, including apoptosis, tumorigenesis, neuronal differentiation, stress responses, and cell-cycle regulation (27–29). Based on structural conservation of the BAG domain, 7 BAG-like proteins can be identified in *Arabidopsis thaliana* (Arabidopsis) (30). These proteins are classified into two groups according to their structural features. The first group, BAG1-BAG4, contains both a BAG domain and a ubiquitin-like (UBL) domain, whereas the second group, BAG5-BAG7, is characterized by a unique calmodulin (CaM)-binding motif located near the BAG domain. Among these proteins, BAG4 has been implicated in responses to various environmental stresses, such as drought, salinity, UV irradiation, and oxidative stress (30). However, the molecular mechanisms underlying BAG4-mediated stress responses remain poorly understood.

In this study, we found that, in addition to BAG4, BAG1-BAG3 also associate with AP-2 and TPC. Although the *bag1bag2bag3bag4* quadruple mutant exhibited altered vegetative growth, it did not show pronounced defects in endocytic recruitment or dynamics. Instead, it displayed increased sensitivity to salt stress and multiple cellulose biosynthesis inhibitors. BAG1-BAG4 interacted with CESA6, and salt-induced CESA6 ubiquitination and degradation were enhanced in the quadruple mutant. Together, these findings suggest that BAG1-BAG4 function redundantly to maintain CESA6 stability under salt stress.

## Results

### BAG1-BAG4 are required for Arabidopsis growth

We previously identified the chaperone regulator, BAG4 as an interactor of several subunits of the CME adaptor complexes AP-2 and TPC (15). However, the functional significance of these interactions remained unknown. BAG4 shares conserved sequence regions and a similar domain architecture with its close homologs BAG1–BAG3 (Fig. 1*A* and *SI Appendix*, Fig. S1 *A* and *B*), suggesting that they may have overlapping functions. Consistently, co-immunoprecipitation (Co-IP) analysis in *Nicotiana benthamiana* leaf epidermal cells or using transgenic Arabidopsis lines expressing BAG-GFP fusion proteins revealed interaction of AP2S and TPLATE with all four BAG proteins (*SI Appendix*, Fig. S1*C* and Fig. 1*B*). Together, these results indicate that BAG1, BAG2, BAG3, and BAG4, can each associate with the core endocytic machinery.

**Fig. 1.**
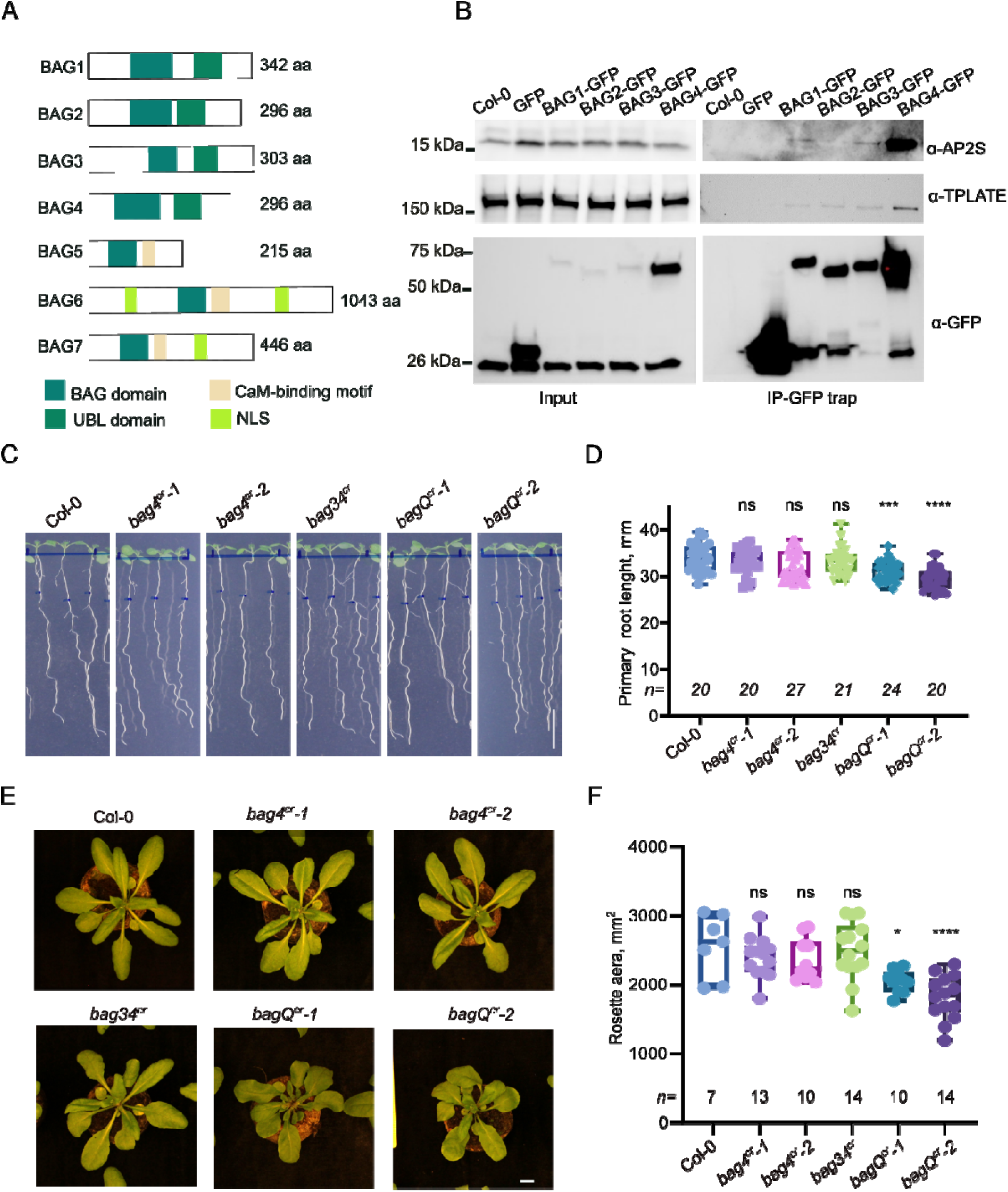
BAG1-BAG4 are required for Arabidopsis growth. (A) Schematic representation of the Arabidopsis BAG-family proteins, indicating protein length (aa: amino acids) and the positioning of various protein domains. UBL, ubiquitin-like domain; NLS, nuclear localization signal; CaM, calmodulin-binding motif. (B) Co-immunoprecipitation (co-IP) assays in Arabidopsis. Protein extracts from plants (*bagQ^cr^-2* mutant) expressing *pRPS5a-gBAGs-GFP* were subjected to pull-down using GFP-trap; AP2S and TPLATE were detected with specific antibodies. Wild-type (Col-0) and *p35S-GFP-*expressing Arabidopsis plants were used as negative controls. (C) Wild-type and *bag* mutants were germinated and grown for 5 days on agar plates and then transferred to new agar media and grown for 4 additional days. Scale bar, 1 cm. (D) Primary root length quantification of seedlings in (C). (E) Wild-type and *bag* mutants were germinated and grown for 7 days on agar plates and then transferred to soil and grown for 3 additional weeks. Scale bar, 1 cm. (F) Rosette area quantification of soil-grown plants in (E). Box plots show the median, 25^th^–75^th^ percentiles, and minimum and maximum values. All individual data points are shown. *n*, number of plants analyzed. **** *P* ≤ 0.0001, *** *P* ≤ 0.001, * *P* ≤ 0.05 ns, not significant. Statistical significance was determined by one-way ANOVA followed by Dunnett’s multiple-comparison test, with each line compared with the wild-type.

To investigate the role of the four BAG proteins, a quadruple mutant was generated using multiplex CRISPR. Two quadruple mutants with different allele combinations were identified: *bag1^cr^-1bag2^cr^-1bag3^cr^-1bag4^cr^-1* (*bagQ^cr^-1*) and *bag1^cr^-2bag2^cr^-2bag3^cr^-1bag4^cr^-1* (*bagQ^cr^-2*) (*SI Appendix*, Fig. S2). When grown *in vitro* for 9 days, the primary roots of the *bagQ^cr^*mutants were 10% shorter than those of wild-type, representing a significant difference from the wild-type (Fig. 1 *C* and *D*). In contrast, the *bag4^cr^-1* single and *bag3^cr^-1bag4^cr^-1(bag34^cr^)* double mutants did not exhibit any notable phenotypic changes compared to wild-type. After four weeks of growth in Jiffy pellets, the *bagQ^cr^*mutants displayed a compact growth and an approximately 25% reduction in rosette size compared with wild-type plants, whereas no such phenotypic change was observed in the *bag4^cr^-1* single and *bag34^cr^*double mutants (Fig. 1 *E* and *F*). These observations suggest that multiple BAG proteins are required for Arabidopsis growth under standard growth conditions.

### BAG proteins are not core regulators of CME

To assess whether the four BAG proteins influence CME, FM4-64 uptake assays were performed to monitor bulk endocytic flux in plant cells. FM4-64 internalization was reduced in the *bagQ^cr^* mutant compared with the wild-type (*SI Appendix*, Fig. S3 *A* and *B*), indicating reduced endocytosis. However, the protein amounts of AP2S and TPLATE subunits were not affected in *bagQ^cr^* mutants (*SI Appendix*, Fig. S3*C*). To further investigate whether the reduced internalization was caused by altered PM recruitment of AP-2 or TPCs, *pAP2M-AP2M-GFP* and *pLAT52-TPLATE-GFP* constructs were transformed into the *bagQ^cr^-2* mutant. One transgenic line for each construct was backcrossed to Col-0, and the F1 heterozygous plants were used as a control. The cytoplasm-to-PM fluorescence ratio of TPLATE-GFP and AP2M-GFP was comparable between the *bagQ^cr^*mutant and the heterozygous plants, indicating that loss of the four BAG proteins did not interfere with the PM recruitment of the endocytic machinery (*SI Appendix*, Fig. S3 *D-G*). TPLATE-GFP lifetimes varied across multiple independent *bagQ^cr^-2* and wild-type lines. Despite a trend toward shorter lifetimes in the mutants, this variability did not support a robust genotype-dependent effect (*SI Appendix*, Fig. S3*H-I*). Taken together, these results suggest that BAG deficiency affects endocytic flux without substantially altering AP-2 or TPC recruitment to the PM or detectably affecting TPC dynamics.

### *bagQ^cr^* mutants are hypersensitive to salt stress

BAG4 has previously been implicated in plant responses to salt stress (30). To examine the sensitivity of the *bag* mutants to salt, 5-day-old seedlings were transferred to medium containing 110 mM NaCl and grown for an additional seven days before primary root length was assessed. *bag4^cr^*single and *bag34^cr^* double mutants displayed levels of root growth inhibition comparable to those of the wild-type (Col-0). However, the two *bagQ^cr^* mutants exhibited severe root-growth arrest, swollen root tips and extensive root hair formation (Fig. 2 *A* and *B*). The salt-hypersensitive phenotype of *bagQ^cr^-2* mutant was partially complemented by introducing a genomic construct of BAG4 as well as by GFP-fusion of BAG1, BAG2 and BAG3, but not by GFP-fused versions of BAG4 (Fig. 2 *C* and *D* and *SI Appendix*, Fig. S4*A*). BAG1-GFP, BAG2-GFP, BAG3-GFP, BAG4-GFP and GFP-BAG4 localized predominantly to the cytoplasm and nucleus (*SI Appendix*, Fig. S4*A-B,* Movie S1), consistent with previous reports (28). However, BAG1, BAG2, and BAG3, but not BAG4 also localized to the cell plate and at the PM (*SI Appendix*, Fig. S4*B*). In etiolated hypocotyls, BAG1-GFP, BAG2-GFP, and BAG3-GFP were enriched a the PM, whereas BAG4-GFP and GFP-BAG4 remained cytoplasmic (*SI Appendix*, Fig. S4*B*). Together with the inability of either tagged BAG4 construct to complement *bagQ^cr^-2* mutant phenotype, these observations suggest that BAG4, in contrast to the other isoforms, does not tolerate either N- or C-terminal GFP tagging. Collectively, these findings support overlapping functions of BAG1-BAG4 in maintaining Arabidopsis growth under salt stress.

**Fig. 2.**
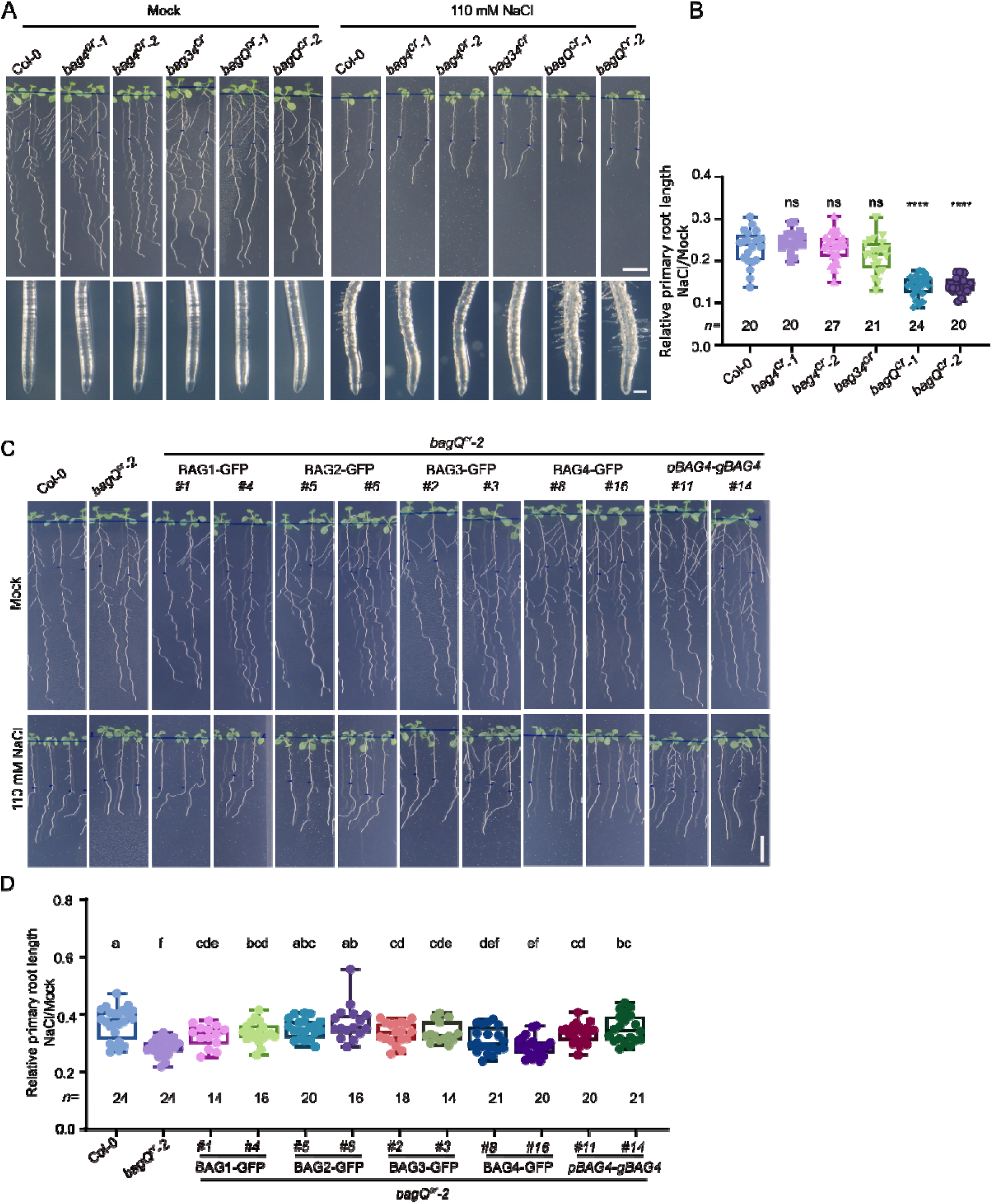
*bagQ^cr^* mutant is hypersensitive to salt stress. (A) Wild-type (Col-0) and *bag* mutants were germinated and grown for 5 days on agar plates and then transferred to medium containing either 110 mM NaCl or mock and grown for additional 7 days (top). Magnified images show root tip swelling in *bag* mutants under salt stress (bottom). Scale bars, 1 cm (top) and 200 μm (bottom). (B) Primary root length quantification of seedlings in (A). (C) Wild-type, *bagQ^cr^-2*, and two independent complemented lines expressing each BAG-GFP fusion protein from the *pRPS5a* promoter, except for BAG4, which was expressed from its native promoter. Seedlings were germinated and grown as in (A). (D) Primary root length quantification of seedlings in (C). Box plots (B and D) show the median, 25^th^–75^th^ percentiles, and minimum and maximum values. All individual data points are shown. *n*, number of plants analyzed. **** *P* ≤ 0.0001, ** *P* ≤ 0.01, ns, not significant. The same letter indicates no significant difference, whereas different letters indicate a significant difference (P ≤ 0.05). Statistical significance was determined by one-way ANOVA followed by Dunnett’s (B) or Tukey’s (D) multiple-comparison test, with each line compared with Col-0 (B).

### *bagQ^cr^* mutants are hypersensitive to cellulose biosynthesis inhibitors (CBIs)

The previously identified BAG4 interactome (15) contains the TETRATRICOPEPTIDE THIOREDOXIN-LIKE (TTL1) and TTL3 proteins, which mediate salt-stress responses in Arabidopsis by regulating the stability of CESA6 (32). We therefore hypothesized that BAG proteins might also affect cellulose biosynthesis. To test this possibility, we examined the responses of *bag* mutants to three CBIs: C17 (33), isoxaben (23) and Endosidin20-1 (ES20-1) (34). Whereas the *bag4^cr^* single and the *bag34^cr^* double mutants responded similarly to wild-type plant under all CBI treatments, the *bagQ^cr^* mutants exhibited pronounced hypersensitivity to these inhibitors (Fig. 3 *A-D* and *SI Appendix*, Fig. S4 *C-E*), characterized by severe root growth arrest, root tip swelling and ectopic root hair formation, consistent with impaired cellulose biosynthesis (Fig. 3 *A-D* and *SI Appendix*, Fig. S4 *C-E*). Likewise, when grown in darkness, *bagQ^cr^-1* and *bagQ^cr^-2* seedlings developed significantly shorter and thicker hypocotyls than wild-type, *bag4^cr^*single and *bag34^cr^* double mutants (Fig. 3 *D-F*). Together, these findings suggest that BAG1-BAG4 function in maintaining cellulose biosynthesis and that defects in this process may contribute to the impaired salt-stress tolerance of *bagQ^cr^* mutants.

**Fig. 3.**
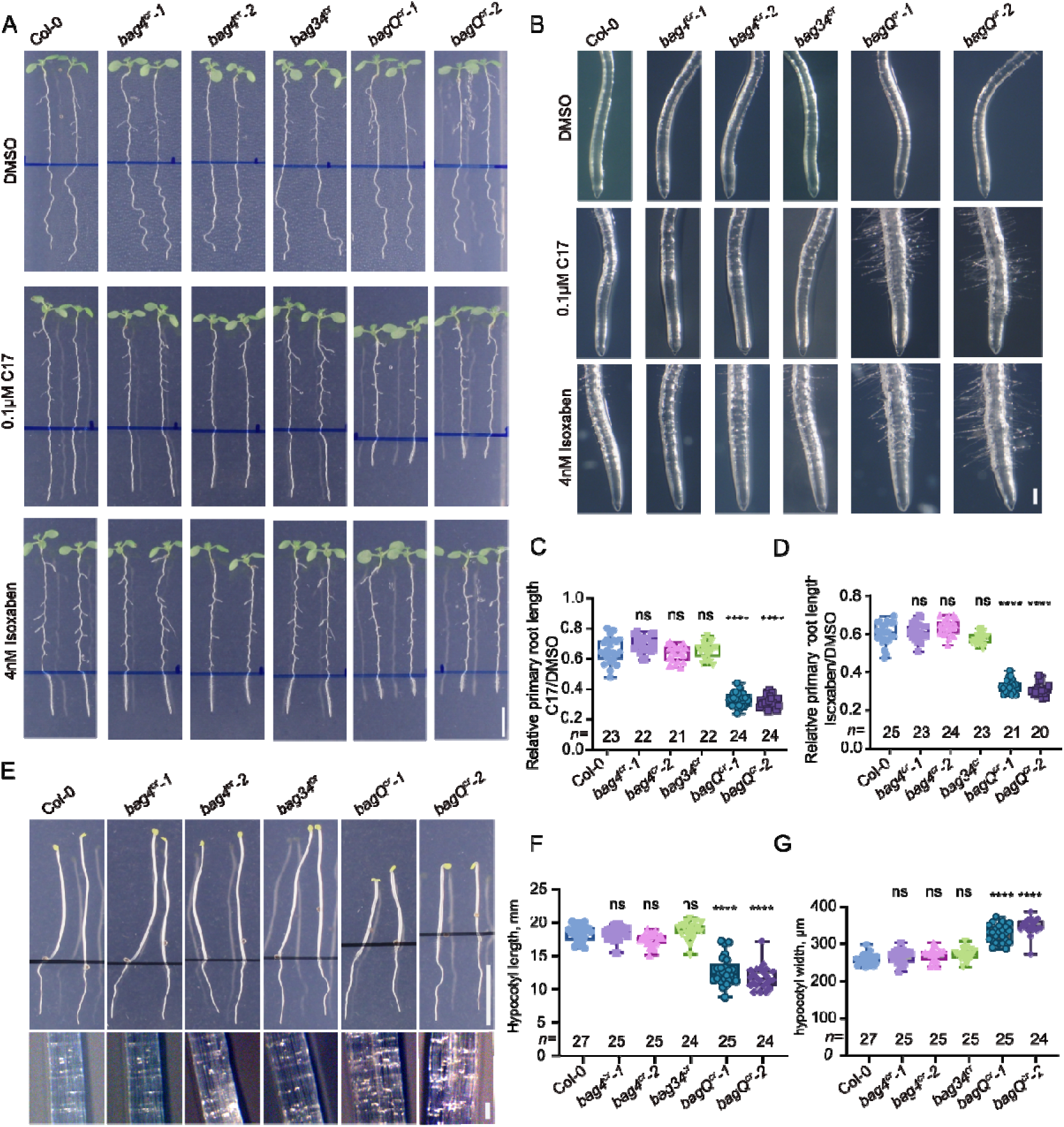
*bagQ^cr^*mutants are hypersensitive to cellulose biosynthesis inhibitors. (A) Wild-type (Col-0) and *bag* mutant seedlings were germinated and grown for 5 days on agar plates, transferred to control media (DMSO) or media supplemented with 0.1 μM C17 or 4nM Isoxaben, and grown for 3 additional days. (B) Magnified images showing root-tip swelling in *bagQ^cr^* mutants following treatment. Scale bars, 1 cm (A) and 200 μm (B). (C and D) Primary root length quantification of seedlings (A). (E) Wild-type and *bag* mutants seedlings germinated for 2 days, transferred to new plates, and grown for additional 5 days in darkness (top). Magnified images show the hypocotyl width (bottom). Scale bars, 10 mm (top) and 200 μm (bottom). (F and G) Quantification of hypocotyl width and length of seedlings in (C). Box plots (C, D, F and G) show the median, 25^th^–75^th^ percentiles, and minimum and maximum values. All individual data points are shown. *n* number of plants analyzed. **** *P* ≤ 0.0001, ** *P* ≤ 0.01, ns, not significant. Statistical significance was determined by one-way ANOVA followed by Dunnett’s multiple-comparison test, with each line compared with the Col-0.

### BAG1-BAG4 proteins are required for maintaining CESA6 stability under salt stress

To determine whether BAG proteins interact with CESA6 and regulate its function, we performed Co-IP assays. GFP-CESA6 was co-expressed with mCherry-tagged BAG1-BAG4 in *Nicotiana benthamiana* leaf epidermal cells. All four BAG proteins co-immunoprecipitated with CESA6 (Fig. 4*A*). To investigate whether BAG proteins regulate CESA6 stability under salt stress, *pCESA6-mNeonGreen-CESA6* and *pUBQ-GFP-CESA6* constructs were introduced into wild-type (Col-0) and *bagQ^cr^-2* mutant plants. Five-day-old *mNeonGreen-CESA6/bagQ^cr^-2* seedlings were transferred to medium with or without 110 mM NaCl for 24 hours. qPCR results showed that CESA expression levels although induced by salt were comparable between wild-type and the *bagQ^cr^* mutant (*SI Appendix*, Fig. S5*A*). However, when the mNeonGreen signal was examined in the salt-stressed root meristem cells, a pronounced vacuole mNeonGreen-CESA6 signal was observed in cells of *bagQ^cr^-2* mutant but not in the wild-type, indicating enhanced vacuolar degradation of CESA6 (Fig. 4*B*). Consistent with this observation, a GFP flux assay (32), revealed increased accumulation of free GFP in *bagQ^cr^* mutant compared to the wild-type (Fig. 4*C*) under salt stress, further supporting enhanced vacuolar turnover of CESA6 in the in the absence of BAG1-BAG4.

**Fig. 4.**
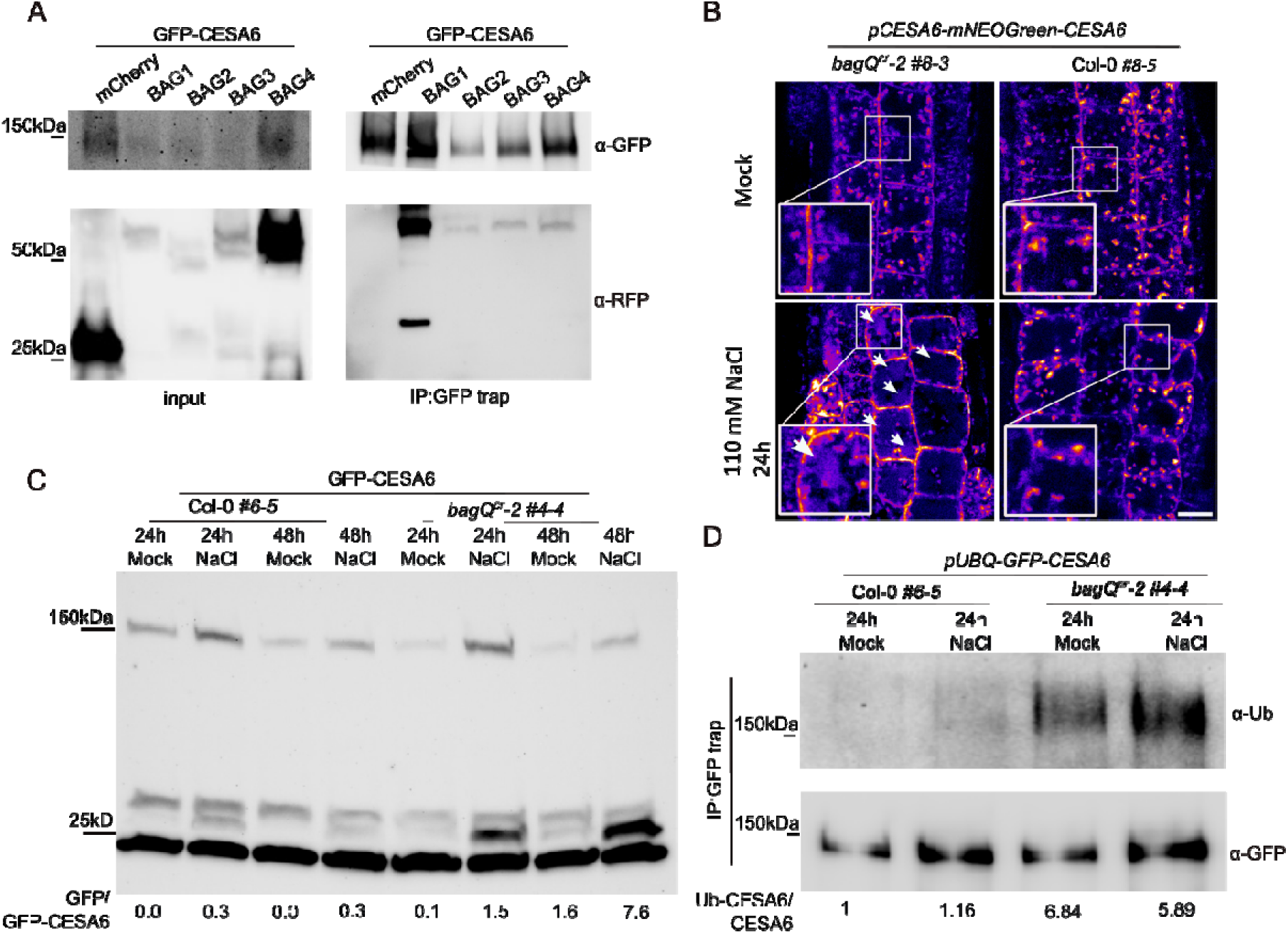
BAG1-BAG4 proteins are required for maintaining CSC stability under salt stress. (A) Co-immunoprecipitation (Co-IP) assay of GFP-CESA6 and mCherry-BAG proteins in *Nicotiana benthamiana* leaves. (B) Localization of mNEOGreen-CESA6 in root epidermal cells of 5-day-old wild-type (Col-0) and *bagQ^cr^-2* seedlings. The arrow and insets indicate the vacuolar localization of CESA6 in the seedlings. (C) Release of free GFP from GFP-CESA6 under salt stress. Seedlings expressing *pUBQ-GFP-CESA6* were germinated on agar medium for 4 days and then transferred to medium with or without 110 mM NaCl for 24 or 48 h. Quantification of the free GFP/GFP-CESA6 ratio from densitometric analysis of the immunoblot is shown below. Total protein extracts were analyzed by immunoblotting with anti-GFP antibodies. (D) Ubiquitination of the CESA6 under salt stress. Seedlings expressing *pUBQ-GFP-CESA6* were germinated on agar medium for 5 days and then transferred to medium with or without 110 mM NaCl for 24 h. Quantification of the Ub-CESA6/CESA6 ratio from densitometric analysis of the immunoblot is shown below. Total protein extracts were immunoprecipitated using GFP trap and subsequently analyzed by immunoblotting with anti-GFP and anti-Ubiquitin antibodies.

Given that ubiquitination is a key post-translational signal regulating the trafficking and degradation of PM proteins (36), we next examined whether the enhanced vacuolar turnover of CESA6 in *bagQ^cr^*was associated with altered CESA6 ubiquitination. Immunoprecipitation of GFP-CESA6 followed by immunoblotting with an anti-ubiquitin antibody revealed increased CESA6 ubiquitination in *bagQ^cr^* compared with wild-type under both mock and salt treatment conditions (Fig. 4*D*). To investigate if we could couple the hypersensitivity to CBIs and the increased degradation of CESA, we assessed PM density of CESA6 under normal and salt-stress conditions. Under control conditions, the PM density of CESA6 was significantly lower in *bagQ^cr^* than in wild-type (*SI Appendix*, Fig. S5*B*). Salt treatment reduced the PM CESA6 density in wild-type, whereas no additional reduction was observed in *bagQ^cr^*. Consequently, the difference in CESA6 density between wild-type and *bagQ^cr^* became less apparent following prolonged salt treatment. Fluorescence recovery after photobleaching (FRAP) analysis showed comparable fluorescence recovery of CESA6 in the wild-type and *bagQ^cr^*, suggesting that CESA6 delivery to the PM was not impaired in the mutant (*SI Appendix*, Fig. S5*C*). Hence, the reduced CESA6 density observed in *bagQ^cr^* is unlikely to result from impaired delivery to the PM but may instead reflect enhanced degradation after internalization. Collectively, these results indicate that BAG proteins function together with CESA6 to regulate its stability under salt-stress conditions. Loss of BAG proteins leads to increased CESA6 ubiquitination and enhanced vacuolar turnover.

## Discussion

The Arabidopsis BAG protein family comprises seven members, but functional studies so far have only focused on a few. BAG5 regulates leaf senescence through CaM/Hsc70 (34), BAG6 is required for autophagy and fungal resistance (38), and BAG7 mediates heat and cold tolerance via the unfolded protein response (39, 40). By contrast, the functions of BAG1-BAG4 remain poorly understood. Some studies suggested that BAG4 is involved in biotic and abiotic stress tolerance (30, 41), yet the underlying mechanisms and the potential redundancy among these closely related proteins remain unclear. BAG4 was initially identified in the interactomes of AP-2 and TPC (15), and our subsequent analysis showed that BAG1-BAG3 also associate with components of both complexes. This shared association suggests that BAG1-BAG4 may function within a common protein network linked to CME. Consistent with this, the *bagQ^cr^* mutant showed reduced FM4-64 uptake, indicating that bulk endocytosis is affected when BAG1-BAG4 are lost. However, we could not attribute this phenotype to defective PM recruitment of TPLATE and AP-2 subunits, nor did we detect significant changes in TPLATE PM dynamics. Although a trend toward shorter TPLATE lifetimes was observed, the high variability among independent transgenic lines precluded a clear conclusion. One possibility is that loss of BAG function does not affect TPC and AP-2 membrane recruitment, but instead destabilizes endocytic progression, thereby resulting in shorter lifetimes at the PM. Such changes may be difficult to detect because automated quantification is inherently prone to generating very short lifetime measurements, potentially masking a shift toward shorter events. By contrast, delayed CME caused by prolonged retention of the endocytic machinery at the PM results in longer lifetimes, which are more readily detected (36, 42, 43). Previous studies have shown that AP-2 and TPC cooperate in CESA internalization, with impairment of either machinery resulting in increased CESA density at the PM (7–9). In contrast, CESA density was reduced in the *bagQ^c^*^r^ mutant, which may result from enhanced CESA destabilization in the absence of BAG1-BAG4 proteins. The localization of BAG1-BAG4 proteins and their interaction with the CME machinery point to a role for BAG1-BAG4 at the PM. However, the enhanced CESA destabilization in *bagQ*^cr^ is not reflected by increased endocytosis, as endocytic flux was reduced in the mutant. Although the reduced FM4-64 uptake could be interpreted as a secondary consequence of impaired cellulose biosynthesis (44, 45), this was not accompanied by the expected increase in TPC lifetime at the PM. It therefore remains unclear at which endosomal compartment BAG proteins exert their stabilizing effect on CESA.

Notably, *bagQ^cr^* displays several hallmarks of cellulose-related defects, including isotropic growth under salt and dark conditions and hypersensitivity to CBIs (23, 24, 32). Further supporting a link to cellulose biosynthesis, BAG1-BAG4 interacted with CESA6, and salt-induced ubiquitination and degradation of CESA6 was enhanced in the *bagQ^cr^* mutant. How BAG1-BAG4 maintain CESA6 stability remains an open question. BAG proteins are often associated with protein degradation through quality control and chaperone-related processes under stress (25, 27, 46). Interestingly, here BAGs act as a CESA proteostasis factors rather than CESA degradation factors. One possibility is that BAGs help stabilize CESA6 directly or indirectly by regulating its folding, assembly, or degradation under salt stress. Another possibility is that BAGs act at the interface between trafficking and proteostasis, influencing whether CESA6 is recycled, retained, or degraded. Loss of BAG shifts CESA toward degradation, reducing the amount of functional cellulose synthase complexes at the PM. Further work will be required to determine the exact mechanism of BAG regulation of CESA6 degradation. It is interesting to highlight that although structurally similar, BAG4 in contrast to BAG1-BAG3 did not tolerate GFP-tagging. This may reflect BAG4’s broader interactome, including nuclear and stress-related proteins (15). Therefore, although BAG1-BAG4 might redundantly regulate cellulose biosynthesis and stress responses, individual members likely also contribute to other pathways. For example, rice OsBAG4 has been reported to modulate potassium transporter expression (47, 48).

In summary, we reveal a previously unrecognized role for BAG1-BAG4 in maintaining cellulose synthase stability during salt stress. We show that BAG1-BAG4 function redundantly to support CESA6 homeostasis, thereby linking BAG-mediated stress protection to cellulose synthesis and cell wall integrity. Our findings establish a role for plant BAG proteins in maintaining cell wall homeostasis under stress.

## Materials and Methods

### Plant material, growth conditions and chemical treatments

*Arabidopsis thaliana* (L.) Heynh., accession Columbia-0, plants were used for all experiments. Arabidopsis seeds were sown on half-strength Murashige and Skoog (½MS) agar plates with 1% sucrose and vernalized at 4°C under dark conditions for 3 days. Seeds were germinated and grown at 22°C and a 16-h light/8-h dark photoperiod for 5, 7 or 9 days, according to the experimental purposes. For the salt or chemical treatment assay, seeds were germinated and grown on ½MS for 3 or 5 days then transformed to the ½ MS plates with or without 110mM NaCl, 4nM Isoxaben, 2 µM C17, 1 µM Endosin20-1 or DMSO and a 16-h light/8-h dark photoperiod for 5, 7 or 9 days. NaCl (AnalytiChem, 7647-14-5), Isoxaben (Sigma-Aldrich, Catalogue Number: 36137), C17 (100 uM stock in DMSO) (33), ES20-1 (1 mM stock in DMSO) (34) were used at the concentrations indicated. Transgenic Arabidopsis lines expressing *p35S-GFP/Col-0* (15), *pAP2M-AP2M-GFP/ap2m-2* (49), *pLAT52-TPLATE-GFP/tplate* (22), and *ap2m-2* (49) have been previously described. *Nicotiana benthamiana* plants were grown in the greenhouse under a normal 14-h light/10-h dark regime at 25°C.

### Rosette area and root length measurements

Four-week-old plants grown in jiffy pellet in multipot trays were imaged. The total leaf area of individual plants was selected by means of the ImageJ color threshold tool. Per genotype, 7 to 14 seedlings were planted randomly to exclude position effects. Primary root length was measured at the indicated time point after germination. Plates were photographed at a fixed distance with a ruler included for calibration. Root length was quantified from the root to the mark line using ImageJ. The segmented-line tool was used to trace the primary root when necessary, and measurements were converted to millimeters based on the image scale. Root length data were collected from the indicated number of independent seedlings for each treatment and used for subsequent statistical analysis.

#### Generation of constructs and transformation

For the CRISPR/Cas9 constructs, gRNAs that targeted the exons of BAG1, BAG2, BAG3, BAG4 were designed with the ‘CCtop’ software (https://cctop.cos.uni-heidelberg.de:8043/) (50). For the BAG4-Cas9 vector, the two gRNAs of BAG4 were cloned into the entry vectors *pGG-A-AtU6-26-BbsI-ccdB-BbsI-B* and *pGG-B-AtU6-26-BbsI-ccdB-BbsI-C* according to the previously described protocol (15). The CRISPR/Cas9 expression construct was generated by assembling the entry clones with *pGG-C-linker-G*, into the backbone vector *pFASTGK-Atcas9-AG* with the Golden Gate reaction and transformed to the Col-0. For the BAG1,2,3-Cas9 vector, gRNA1 and gRNA2 of BAG1, BAG2 and BAG3 genes were cloned into *pGG-A-AtU6-26-BbsI-ccdB-BbsI-B, pGG-B-AtU6-26-BbsI-ccdB-BbsI-C, pGG-C-AtU6-26-BbsI-ccdB-BbsI-D, pGG-D-AtU6-26-BbsI-ccdB-BbsI-E, pGG-E-AtU6-26-BbsI-ccdB-BbsI-F and pGG-F-AtU6-26-BbsI-ccdB-BbsI-G,* then these entry clones were assembled into the *pFASTGK-Atcas9-AG* with Golden Gate reaction and transformed to the *bag4^cr^-1*. For the BAG1,2-Cas9 vector, gRNA3, gRNA4 and gRNA5 of BAG1 and BAG2 genes were cloned into *pGG-A-AtU6-26-BbsI-ccdB-BbsI-B, pGG-B-AtU6-26-BbsI-ccdB-BbsI-C, pGG-C-AtU6-26-BbsI-ccdB-BbsI-D, pGG-D-AtU6-26-BbsI-ccdB-BbsI-E, pGG-E-AtU6-26-BbsI-ccdB-BbsI-F and pGG-F-AtU6-26-BbsI-ccdB-BbsI-G,* then these entry clones were assembled into the *pFASTGK-Atcas9-AG* with Golden Gate reaction and transformed to the *bag34^cr^*. For genotyping, the genomic DNA sequence of the transgenic plants were amplified by PCR with the primers listed in the Supplementary.

For the *p35S* or *pRPS5a* promoter driven expression vectors of BAG family proteins. Genomic DNA of Arabidopsis BAG1, BAG2, BAG3 and BAG4, were cloned into *pGGB000* and subcloned into the backbone vector pFASTRK-AG by the Golden Gate system *with pGG-A-35SP-B or pGG-A-pRPS5a-B, pGG-C-GFP-D/pGG-C-mCherry-D and pGG-D-35ST-G.* The vectors were finally transformed into Col-0 or the *bag* mutants. For *pBAG4-gBAG4*, a 1,762-bp promoter fragment upstream of the BAG4 start codon was amplified together with the *BAG4* genomic sequence and cloned into *pGGB000*. The resulting construct was subcloned into the backbone vector *pFASTRK-AG* using the Golden Gate system together with *pGG-A-linker-B, pGG-C-linker-D,* and *pGG-D-35ST-G.* For *pBAG4-GFP-BAG4*, a 1,762-bp promoter fragment upstream of the *BAG4* start codon was amplified and cloned into pGGA000. The resulting construct was then subcloned into pFASTRK-AG using the Golden Gate system together with *pGG-B-GFP-C, pGG-C-gBAG4-D,* and *pGG-D-35ST-G*.

### Quantitative RT-PCR

RNA was extracted from 9-day-old seedlings with the ReliaPrep™ RNA Miniprep Systems (Promega). Of purified RNA, 1 μg was amplified in a reverse transcriptase reaction with the qScript XLT 1-Step RT-PCR Kit (Quantabio). Subsequently, qPCR was run with the SYBR Green master mix (Roche) with gene-specific primers designed to amplify. *ACTIN2* was used as the normalization reference. Primers are listed in the Supplementary.

### Co-IP assay

For co-IP in Arabidopsis, seeds were germinated and grown on ½ MS. Seven-day-old seedlings were harvested and ground into powder with liquid nitrogen. The fine powder was resuspended in extraction buffer (50 mM Tris-HCl, pH 7.5, 150 mM NaCl, 0.5% [v/v] NP-40, and complete EDTA-free protease inhibitor cocktail [Roche]). Seeds were germinated and grown on ½MS. Seven-day-old seedlings were harvested and ground into powder with liquid nitrogen. The fine powder was resuspended in extraction buffer (50 mM Tris-HCl, pH 7.5, 150 mM NaCl, 0.5% [v/v] NP-40, and complete EDTA-free protease inhibitor cocktail [Roche]).

For co-IP in *Nicotiana benthamiana*, Agrobacterium cultures carrying the indicated constructs were resuspended in infiltration buffer (10 mM MgCl2, 10 mM methyl ethyl sulfide, pH 5.6, 100 µM acetosyringone) and incubated at room temperature for 2 h. The bacterial cultures were adjusted to a final OD600 = 0.5 for each construct and infiltrated into the leaves. After 3 days incubation in the greenhouse, leaves were harvested, ground into powder with liquid nitrogen and then resuspended in extraction buffer (150 mM Tris-HCl, pH 7.5, 150 mM NaCl, 10% [v/v] glycerol, 10 mM EDTA, 1% [v/v] NP-40, and complete EDTA-free protease inhibitor cocktail [Roche]). The extracts were centrifuged at 20,000 × g at 4°C for 15 min. The supernatants were transferred to new 2-ml tubes and centrifuged for an additional 15 min. After centrifugation, the supernatants were incubated with GFP-trap magnetic agarose (Chromotek) at 4°C for 2 h. Beads were washed three times with 1 ml wash buffer (20 mM Tris-HCl, pH 7.5, 150 mM NaCl, 0.25% [v/v] NP-40] and then eluted with 80 µl elution buffer containing 1× NuPAGE™ LDS Sample Buffer (Thermo Fisher Scientific) and 1× NuPAGE® Reducing Agent (Thermo Fisher Scientific) and boiled for 10 min at 90°C.

### SDS-PAGE and immunoblotting

Total protein extracts were obtained by grinding the plant materials into a fine powder with liquid nitrogen. The powder was resuspended in extraction buffer (50 mM Tris-HCl, pH 7.5, 150 mM NaCl, 0.5% [v/v] NP-40, and complete EDTA-free protease inhibitor cocktail [Roche]). The extracts were centrifuged at 20,000 × g at 4°C for 15 min. The supernatants were transferred to new 2-ml tubes and centrifuged for an additional 15 min. The protein extracts were boiled in sample buffer (1× NuPAGE™ LDS sample buffer, 1× NuPAGE® Reducing Agent [Thermo Fisher Scientific]) for 10 min at 90°C and loaded on a 4–20% (w/v) Mini-PROTEAN® TGX™ Precast Protein Gels (Bio-Rad). The proteins were separated by electrophoresis and blotted onto a membrane with trans-Blot Turbo Mini 0.2 µm Nitrocellulose Transfer Packs (Bio-Rad). Membranes were blocked overnight at 4°C in 3% (w/v) skimmed milk dissolved in 25 mM Tris-HCl (pH 8), 150 mM NaCl and 0.05% (v/v) Tween20. The blots were then incubated at room temperature with the monoclonal α-GFP antibody (1:5000) (Miltenyi Biotec, 130-091-833), α-RFP antibody (1:5000) (ChromoTek, 6G6), α-AP2S antibody (1:500) (51), α-tubulin antibody (Sigma-Aldrich, T5168) (1:2000), α-TPLATE antibody (1:500) (42), and α-Ubiquitin antibody (Santa Cruz, sc-8017) (1:2000).

### Fluorescence imaging and FM uptake

All the confocal images were acquired with a Leica SP8X confocal microscope, equipped with a white-light laser operated at 75% of its maximum output and a ×40 water-corrected objective with application of the gating system (0.3-6 ns) for autofluorescence removal, imaging at a resolution of 1024×1024 pixels, 200 Hz speed, 3× line averaging, two-way scan direction. GFP, YFP, RFP and FM4-64 were excited using a white light laser. The excitation and detection window settings were: GFP, 488/500-530 nm; mNEONGreen; 505/515-615 nm; and FM4-64; 515/570-670 nm. For FM4-64 uptake assay, whole 5-day-old seedlings were incubated with 2 μM FM4-64 (Invitrogen, 2 mM stock in water) solution in half-strength MS liquid medium without sucrose at room temperature for 15 min before imaging. The fluorescent PM intensity and intracellular space were measured with ImageJ for the quantification of the FM4-64 uptake and the AP2M-GFP and TPLATE-GFP signals (36, 43, 52). The relative fluorescent cytoplasm-to-PM intensity was calculated. The regions of interest (ROIs) were outlined in individual epidermal cells using the Select Brush and Freehand Selection tools, respectively. Pixel-intensity histograms were then generated for each ROI. Images containing more than 1% saturated pixels were excluded from the analysis. Intensity measurements from the PM and cytoplasm were processed via an in-house excel-based tool to generate the average intensity of the top 100 pixels. These averages were used to generate the ratio values.

### CESA6 dynamics and FRAP assays

For CESA density, microscopy images were taken using Confocal Microscope 3i CSU-W SoRa Spinning Disk (Camera: Andor iXon Life 888 1024×1024 EMCCD). mNEONGreen was excited by laser 488 nm (20%, 400 ms, average 4 times) and observed under a 100X oil lens. Images were processed by Fiji for image analyses using Thunderstorm plugin and data were filtered using the following parameters: photon offset> 0.01, uncertainty between 5 and 60nm, sigma between 5 and 500nm and photon intensity <500000.

To measure fluorescent recovery after photobleaching, microscopy images were taken using Confocal Microscope 3i CSU-W SoRa Spinning Disk (Camera: Andor iXon Life 888 1024×1024 EMCCD). mNEON signal at the plasma membrane was bleached by laser 488 nm (80%, 100 ms) on a round ROI of 14 nm2 of and observed under a 100X oil lens with 488nm laser excitation (20%, 400ms) with 5s intervals between images. Movies were corrected for xy shift using Fast4Dreg plugin.

### Endocytic dynamics imaging and quantification

Endocytic dynamics were imaged using a Nikon Ti microscope equipped with a Perfect Focus System (PFSIII) for Z-drift compensation and a PerkinElmer UltraVIEW spinning-disk system featuring a Yokogawa CSU-X1 scan head, coupled to a Hamamatsu EMCCD C9100-13 camera (512 × 512 pixels). Images of hypocotyl epidermal cells from 4-day-old etiolated seedlings expressing GFP-fusion proteins were acquired using a 100× oil-immersion objective (Plan Apo; NA = 1.45). Excitation was performed at 488 nm, with an emission window between 500 and 550 nm. Time-lapse imaging was conducted for 5 minutes with a 950-ms exposure time at a frame rate of 1 frame per second (fps). Shutter settings were optimized to maximum sample protection while maintaining high scan speeds. Prior to analysis, image sequences were pre-processed in Fiji. To maximize the analyzable area without the need for image rotation, a polygonal region of interest (ROI) was drawn to strictly exclude cell boundaries. The selected area was then isolated using the “Crop” and “Clear Outside” functions in Fiji. The lifetimes of individual endocytic events were measured in MATLAB 2017b using the detection and tracking modules of the cmeAnalysis package, and further processed as previously described (53–55). For this dataset, the analysis script was slightly modified to bypass an error associated with the Filter Gauss2D function. Following the execution of the script, the quality of the analysis was evaluated using an objective, quantitative classification approach. Principal component analysis (PCA) was performed using five parameters extracted from each movie: maximum fluorescence intensity (MAX), average fluorescence intensity (AVG), minimum fluorescence intensity (MIN), standard deviation (sd), and slope of the fluorescence intensity profile (m). The initial classification was based on the manual assessment of each movie, with profiles corresponding to bona fide endocytic events assigned to class 1 and high-fluctuation or otherwise low-quality profiles assigned to class 0. PCA was used to assess the separation of these two predefined groups based on the extracted fluorescence characteristics. Subsequently, linear discriminant analysis (LDA) was performed to determine a quantitative decision boundary separating profiles classified as high- and low-quality. Consistent with visual inspection of the fluorescence intensity profiles, slope and average fluorescence intensity contributed most strongly to the separation between the two groups, reflecting the pronounced fluctuations characteristic of low-quality profiles. Only movies classified as high-quality according to the LDA-based classification were retained for subsequent statistical analyses. This quantitative filtering approach was used to reduce the influence of aberrant fluorescence intensity profiles and the associated bias towards artificially short event lifetimes.

### Statistical analysis

All statistical analyses were done with the GraphPad Prism v.8 software. The multiple comparisons were done by one-way ANOVA with Dunnett’s multiple comparisons test. For statistical analyses in the Fig.S5B was determined using two-way ANOVA followed by Tukey’s test.

## Supporting information

Moive S1

## AI use disclosure

ChatGPT (OpenAI) was used solely to assist with English language editing of the manuscript.

## Acknowledgments

We thank J. Pan for the kind gift of α-AP2S antibodies, L. De Veylder for the kind gift of the C17 compound, M. Botella, and C. Sánchez-Rodríguez for providing published materials. This work was supported by the Research Foundation-Flanders projects (G008416N, G0E5718N and 3G038020 to E.R.), the China Scholarship Council for predoctoral fellowships (to P.W.), the European Research Council (T-Rex project number 682436 to D.V.D. and OMEGA, project number 101043132 to T.B.J), the Research Foundation Flanders PhD fellowship (163826N to N.S.), the Novo Nordisk postdoctoral grant (NNF24OC0094901 to L.C.M.N.), the Novo Nordisk grants (NNF19OC0056076, NNF20OC0060564, NNF0068884, NNF0086341, NNF0084973 to S.P.) and Villum Investigator (25915 to S.P.).

## Author Contributions

P. W., W. S., D. V. D., and E. R. initiated the work. P. W., S. P., D. V. D., and E. R. designed the experiments. P.W. and I. V. performed most of the work; E. M. and N. S. analyzed endocytic dynamics; L. C. M. N., and S. P. analyzed CESA dynamics; Y.D. did modeling; X. Z., T. B. J., S. P., D.V.D., and E.R. provided guidance. P. W., D. V. D., and E. R. wrote the manuscript. All authors commented on the results and the manuscript.

## Supporting Information

**Fig. S1.**
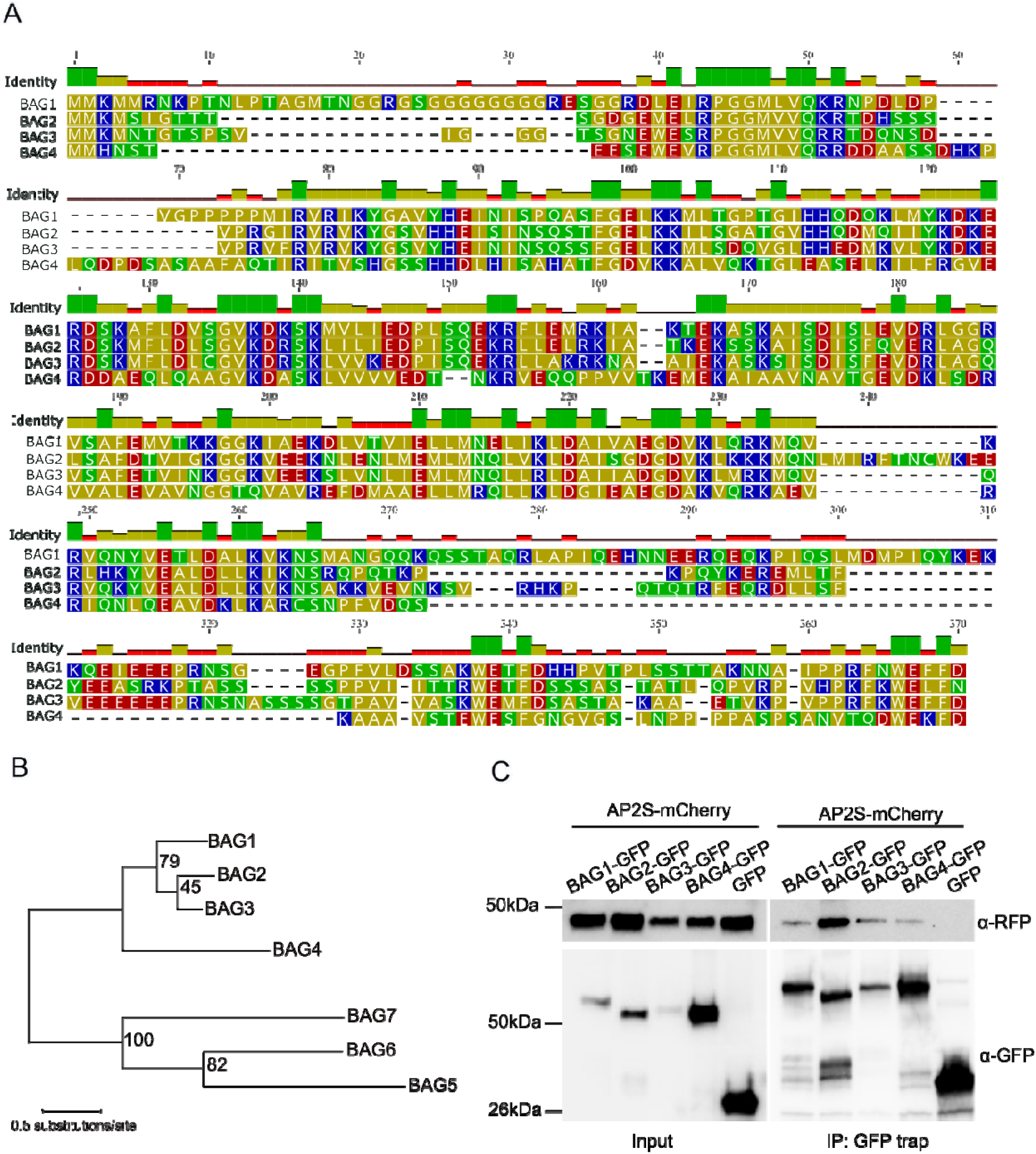
Sequence conservation, phylogeny and interaction of BAG1-BAG4 with AP2S. (A) Multiple sequence alignment of Arabidopsis BAG1-BAG4 proteins, with residues color-coded according to their conservation. (B) Unrooted phylogenetic tree of the BAG family in Arabidopsis. Node values indicate transfer bootstrap expectation (TBE) support based on 100 bootstrap replicates. The scale bar represents 0.5 substitutions per site. (C) Validation of the interactions between AP2S and BAG1-BAG4 proteins by co-immunoprecipitation (co-IP) in *Nicotiana benthamiana* leaves transiently expressing *p35S-AP2S-mCherry*, *p35S-BAGs-GFP* and *p35S-GFP* constructs.

**Fig. S2.**
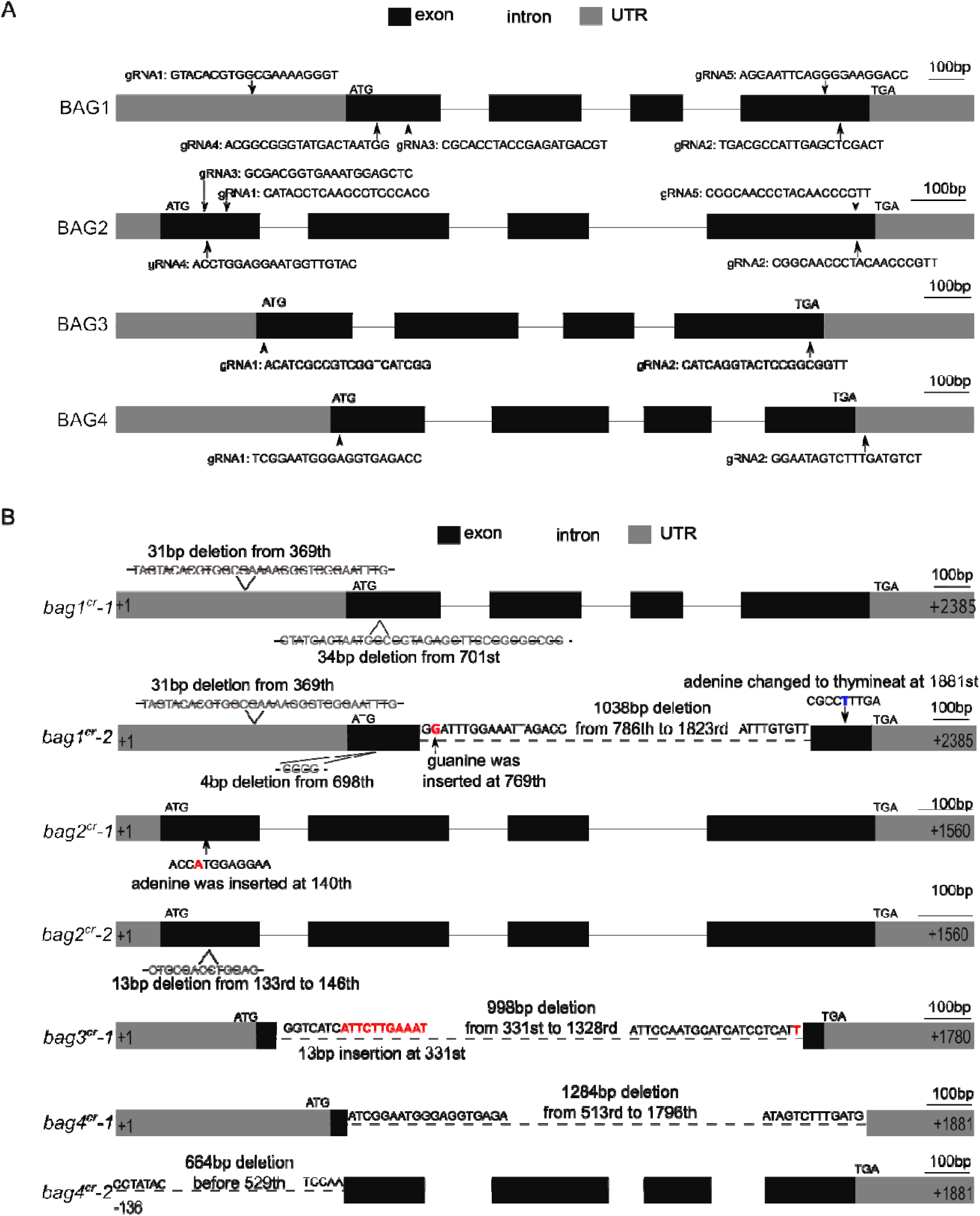
Schematic representation of CRIPSR mutants of the BAG family. A) Schematic representation of the gRNAs targeting *BAG1*, *BAG2*, *BAG3* and *BAG4* genes. B) Schematic representation of the CRISPR-induced edits in the *BAG* genes in the *bag* CRISPR mutants. Red letters, blue letters and dashed lines indicate insertion, nucleotide exchange and deletion, respectively. *bag1^cr^-1* harbored two fragments’ deletions, from base pair (bp) 469 to 499 and from bp 701 to 734, whereas *bag2^cr^-1* contained a single adenine insertion at bp 140 of the BAG2 genomic sequence, resulting in premature termination of protein translation at amino acids 32. The *bag3^cr^-1* carried a 998-bp deletion from bp 331 to the 1328, which removed almost the entire coding region, along with an 11-bp insertion after nucleotide 331. In *bag4^cr^-1*, the genomic region between bp 513 and 1796 was deleted. *bag1^cr^-2* harbored a large deletion from bp 786 to 1823, effectively deleting the entire gene. *bag2^cr^-2* contained a 13-bp deletion after bp 133 of the genomic sequence, leading to premature translation at amino acid the 59. *bag4^cr^-2* contained a 664-bp deletion upstream of bp 528, resulting in the loss of protein translation initiation.

**Fig. S3.**
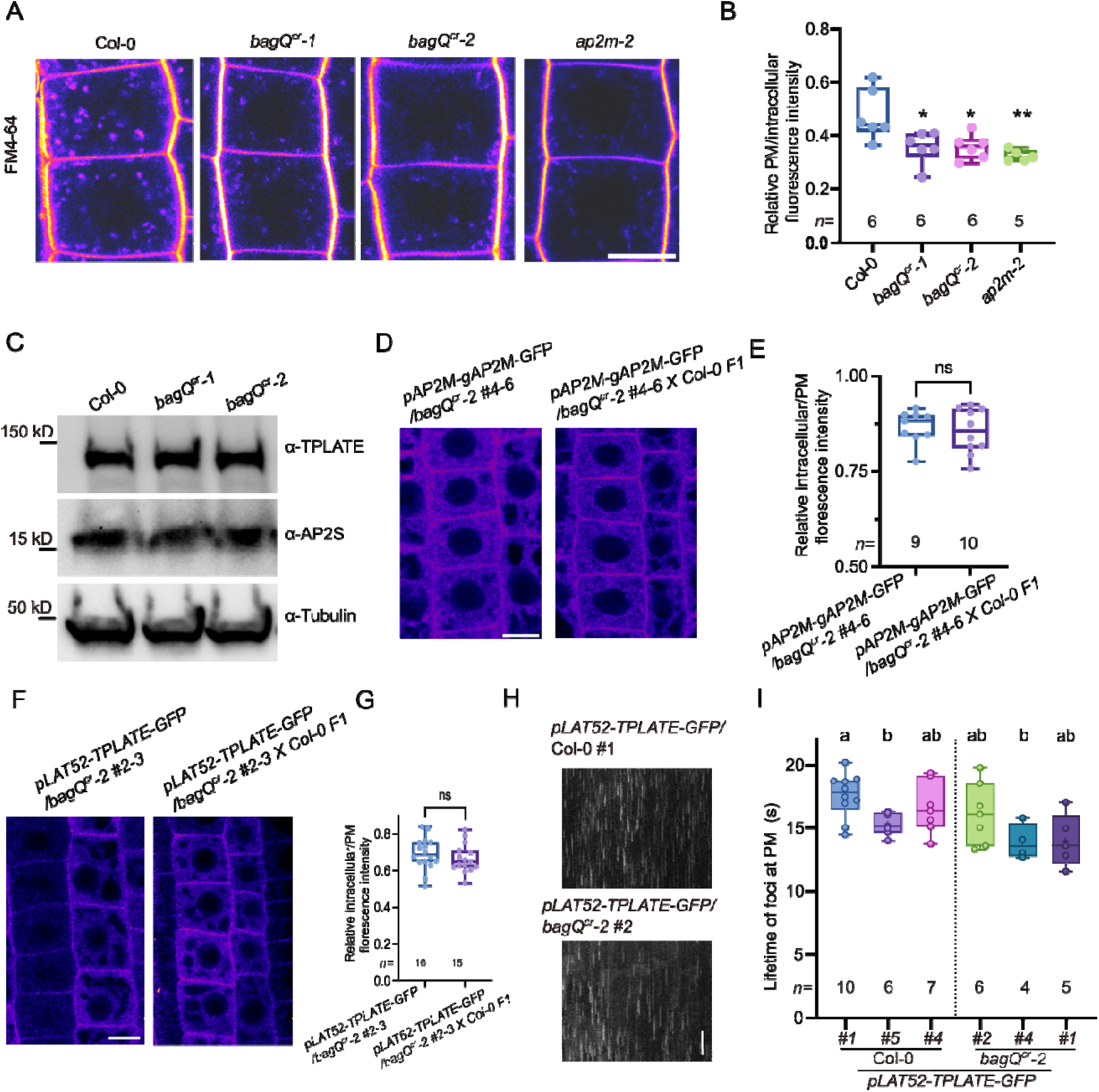
*bagQ^cr^*mutants are affected in bulk endocytosis. (A) FM4-64 uptake in *bagQ^cr^* and *ap2m-2* mutants. Root epidermal cells of 5-day-old seedlings grown on MS medium were imaged after staining with FM4-64 (2 μM, 15 min). (B) Quantification of the ratio of intracellular to plasma membrane (PM) fluorescence intensity in the images in (A). (C) Protein levels of TPLATE, AP2S and tubulin in the wild-type (Col-0) and *bagQ* mutants seedlings. (D and F) Fluorescence intensity images of AP2M-GFP and TPLATE-GFP in root cells of 5-day-old seedlings. (E and G) Ratios of intracellular-to-PM fluorescence intensity of AP2M-GFP and TPLATE-GFP. (H and I) Representative kymographs and quantification of the PM life-time of AP2M-GFP in three independent wild-type and *bagQ^cr^* mutant lines. Scale bars, 10 μm (A, D, F), 60s (H). Box plots (B, E, G and I) show the median, 25^th^–75^th^ percentiles, and minimum and maximum values. All individual data points are shown. *n*, number of plants analyzed. ** *P* ≤ 0.01, * *P* ≤ 0.05, ns, not significant. The same letter indicates no significant difference, whereas different letters indicate a significant difference (*P* ≤ 0.05). Statistical significance was determined by one-way ANOVA followed by Dunnett’s multiple-comparison test, with each line compared with the wild-type (B), compare within all the lines (I), unpaired t-test (E and G).

**Fig. S4.**
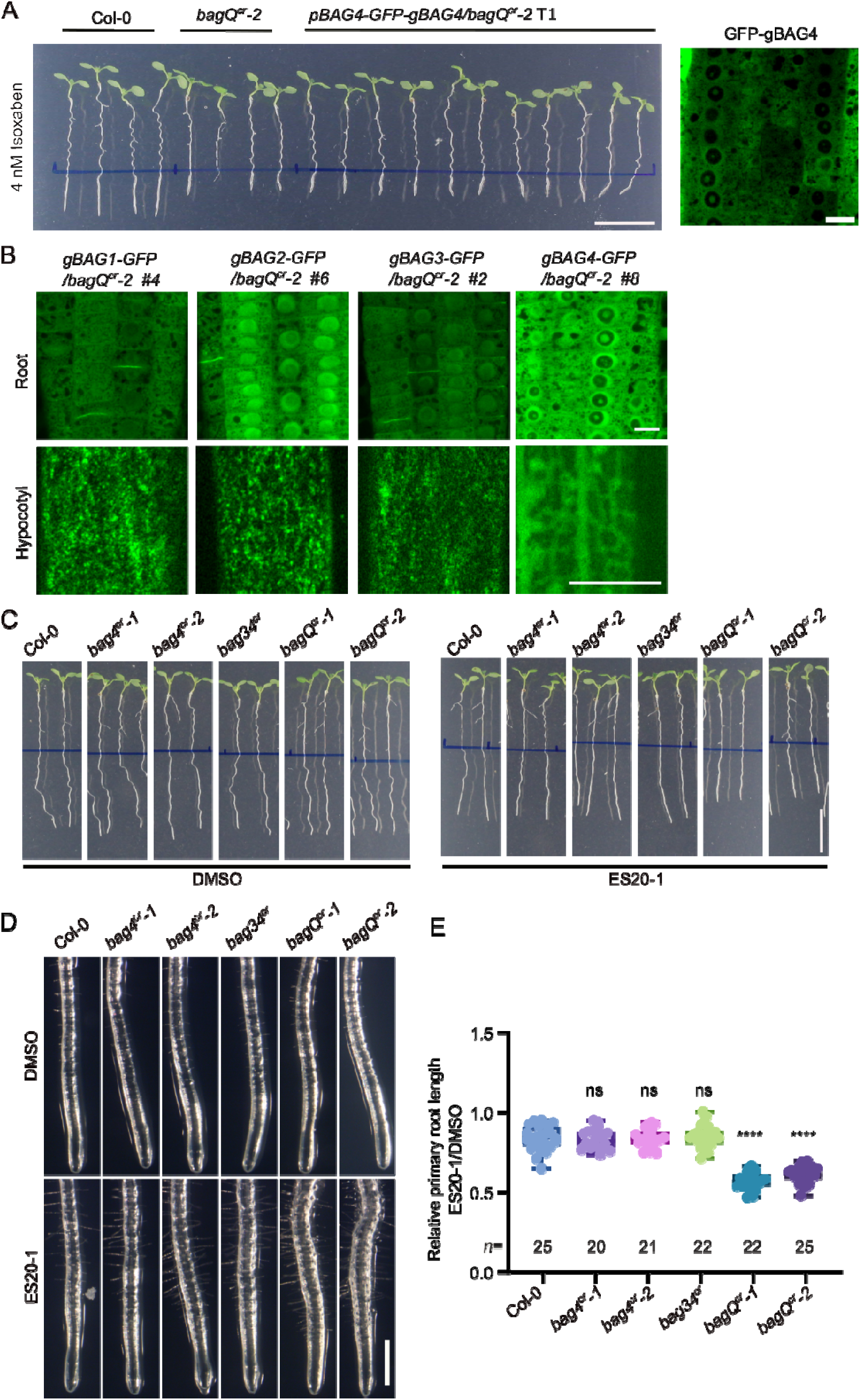
*bagQ^cr^*mutants are hypersensitive to cellulose biosynthesis inhibitors isoxaben and Endosidin20-1 (ES20-1). (A) Wild-type (Col-0), *bagQ^cr^-2*, and *bagQ^cr^-2* transformed with *pBAG4-GFP-gBAG4* construct were germinated and grown for 5 days on MS medium and then transferred to medium containing 4 nM isoxaben for an additional 3 days. Localization of GFP-BAG4 in the root meristem. Scale bars, 1 cm (left), 10 μm (right). (B) Localization of BAG1-GFP, BAG2-GFP, BAG3-GFP and BAG4-GFP in the root meristem (top) and etiolated hypocotyl (bottom) of *bagQ^cr^-2* seedlings expressing each BAG-GFP fusion protein from the *pRPS5a* promoter. Scale bars, 10 μm. (C) Col-0 and *bag* mutant were germinated and grown for 5 days on MS medium and then transferred to control medium (DMSO) or medium supplemented with 1 μM ES20-1 for an additional 3 days. Scale bars, 1 cm (D) Magnified images showing root of untreated (mock) and treated seedlings showing distinctive root-tip swelling in *bagQ^cr^* mutants following ES20-1 treatment. Scale bars, 500 μm. (E) Quantification of primary root length of seedlings in (D). Box plots show the median, 25^th^–75^th^ percentiles, and minimum and maximum values. All individual data points are shown. *n* indicates the number of plants analyzed. **** *P* ≤ 0.0001, ns, not significant. Statistical significance was determined by one-way ANOVA followed by Dunnett’s multiple-comparison test, with each line compared with the wild-type.

**Fig. S5.**
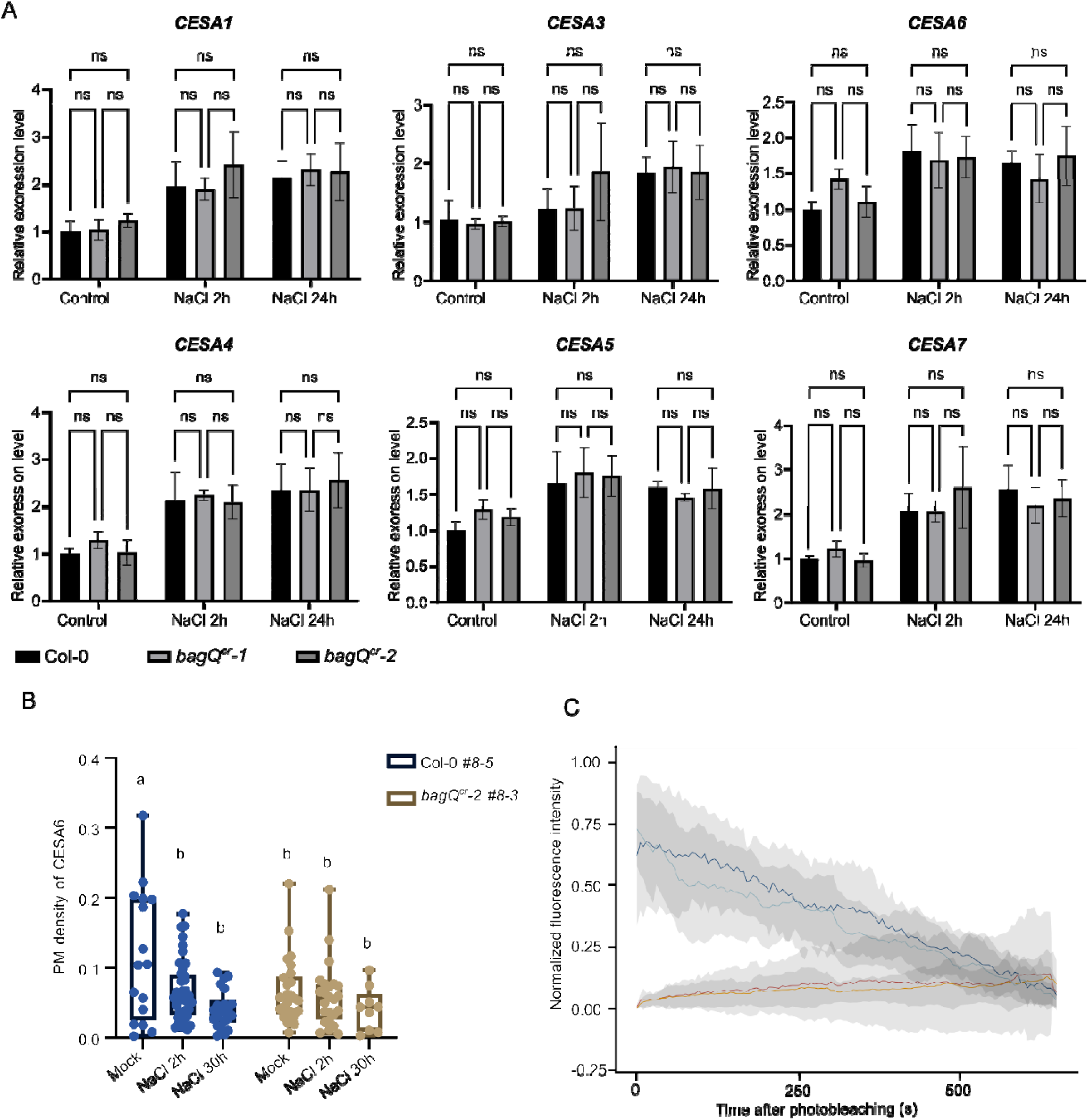
*CESAs* expression and CESA6 dynamics in wild-type (Col-0) and *bagQ^cr^* mutants. (A) Col-0 and *bagQ^cr^* mutants were germinated and grown for 5 days on agar plates and then transferred to the medium supplemented with 110 mM NaCl for 2 or 24 hours (h). Mean ± standard deviation (SD) are shown. ns, not significant. Statistical significance was determined by two-way ANOVA analysis followed by Turkey test. (B) Col-0 and *bagQ^cr^*mutants expressing *pCESA6-mNEON-CESA6* were germinated and grown for 5 days on agar plates and then transferred to the medium supplemented with 110 mM NaCl or mock for 2 or 30 h and the plasma membrane (PM) density of CESA6 was measured. Statistical significance was determined using two-way ANOVA followed by Tukey’s test. The same letters indicate no significant difference, whereas different letters indicate a significant difference (*P* ≤ 0.05). (C) Col-0 and *bagQ^cr^* mutants expressing *pCESA6-mNEON-CESA6* were germinated and grown for 5 days on agar plates before imaging. Mean normalized fluorescence intensity in photobleached ROIs is shown for wild-type (in red) and *bagQ^cr^* (in orange). Mean normalized fluorescence intensity in non-photobleached control ROIs shown for wild-type (in dark blue) and *bagQ^cr^* (in light blue) to account for fluorescence loss during image acquisition. Standard deviation intervals are shown in grey.

**Table S1.**
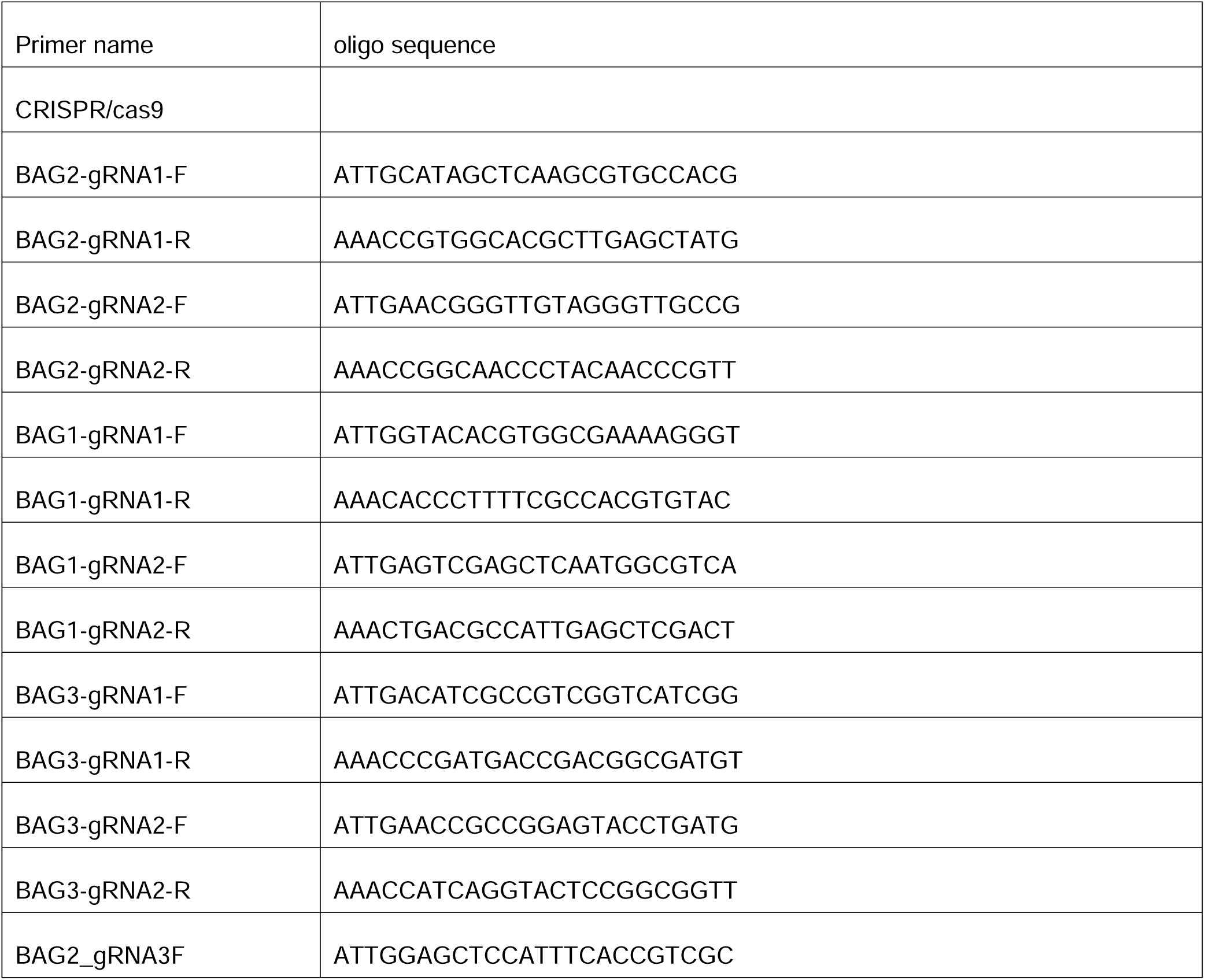
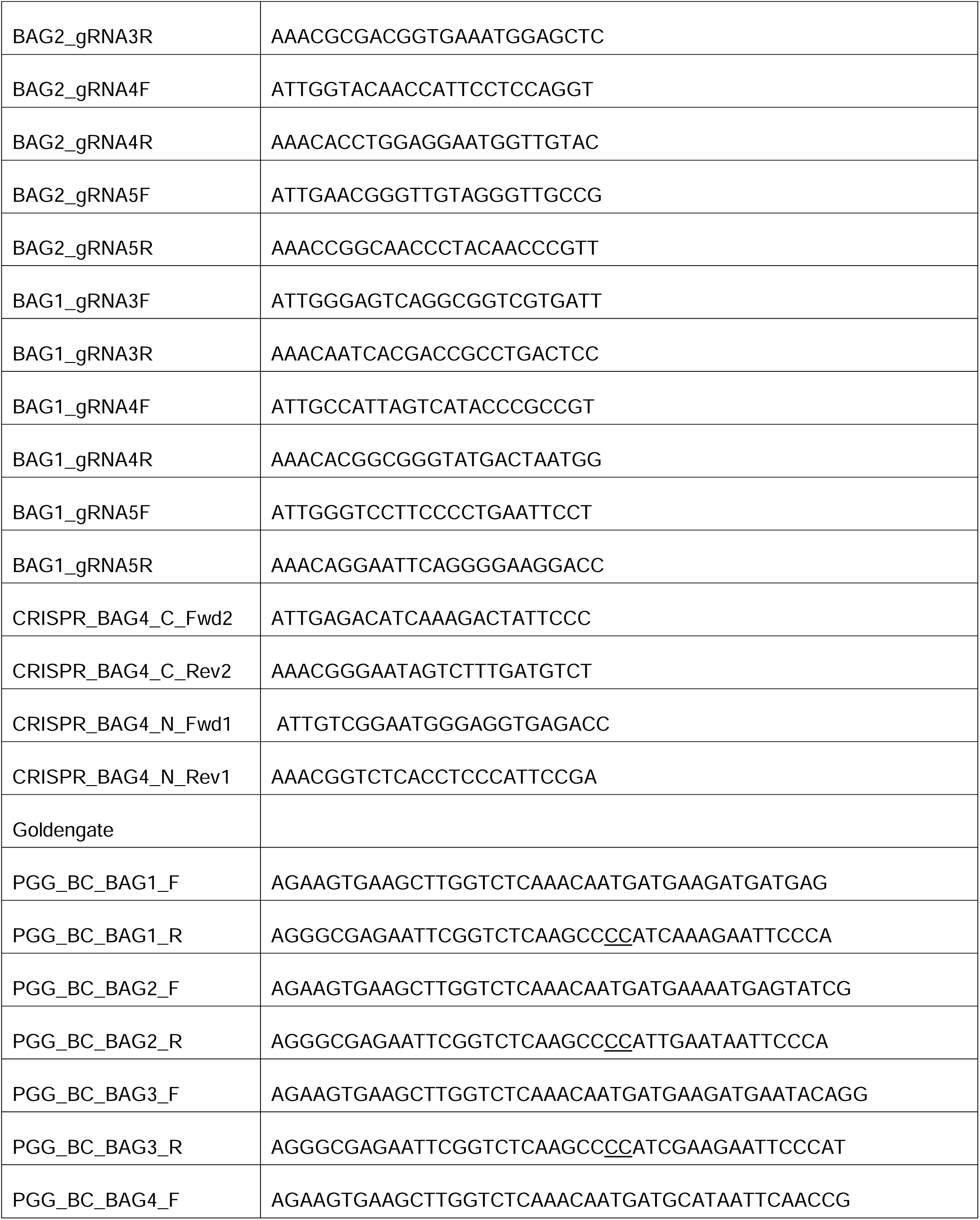
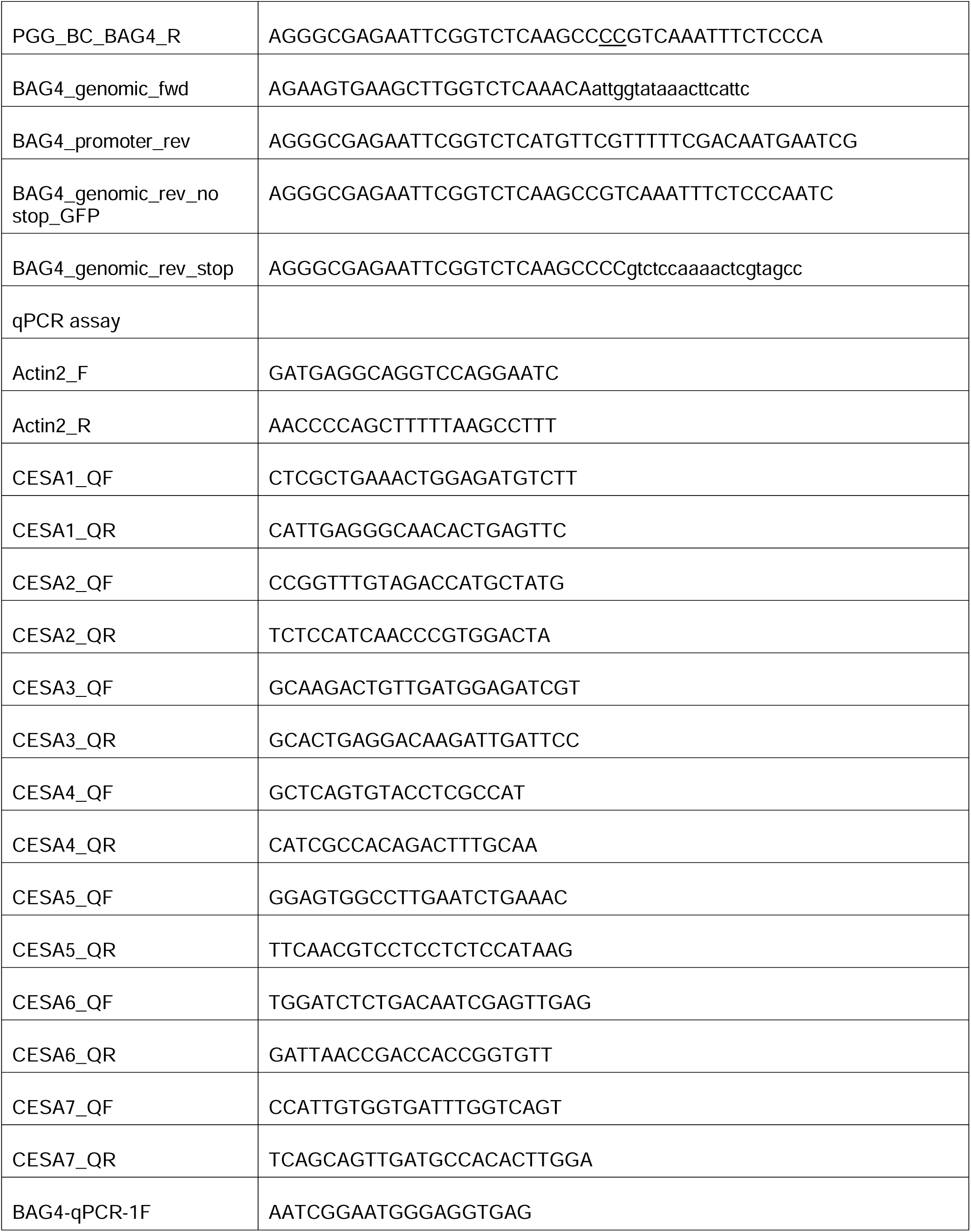
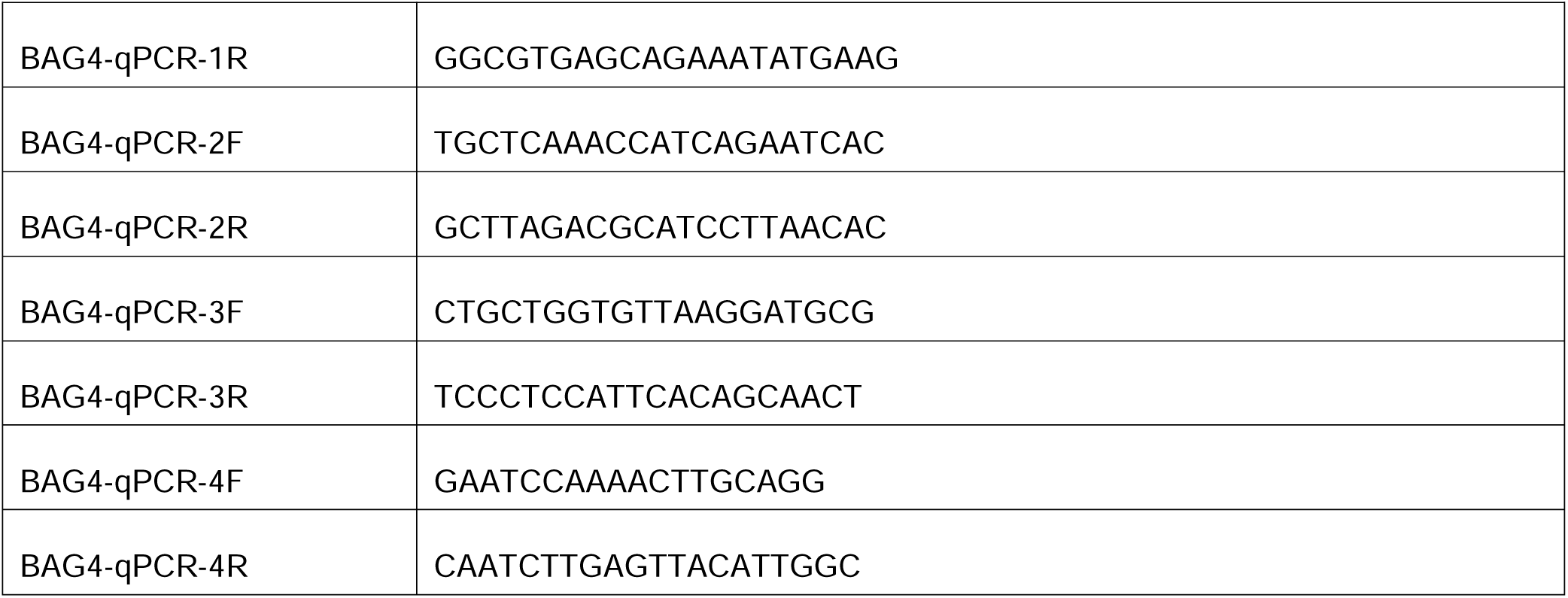
Oligonucleotides used in this study.

**Movie S1. Plasma membrane localization of BAG3-GFP in *bagQ^cr^-2*.** The presented time series were acquired from hypocotyl cells of etiolated Arabidopsis seedlings expressing BAG3-GFP in *bagQ^cr^-2* (BAG3-GFP/*bagQ^cr^-2* #2) using VAEM/spinning-disk confocal microscopy. The movie is shown at a frame rate of seven frames per second (fps).

