## Supplementary figures and images for "Arabidopsis BAG proteins regulate cellulose synthase stability"

### Moive S1

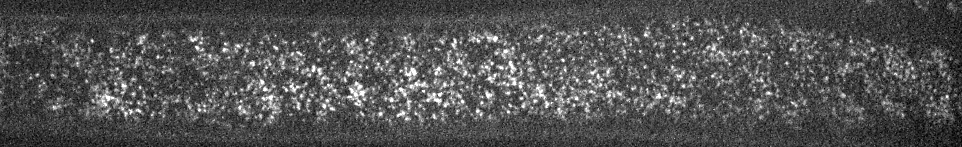
